# Mec1-Independent Activation of the Rad53 Checkpoint Kinase Revealed by Quantitative Analysis of Protein Localization Dynamics

**DOI:** 10.1101/2022.08.04.502543

**Authors:** Brandon Ho, Ethan J. Sanford, Nikko P. Torres, Marcus B. Smolka, Grant W. Brown

## Abstract

The replication checkpoint is essential for accurate DNA replication and repair, and maintenance of genomic integrity when a cell is challenged with genotoxic stress. Several studies have defined the complement of proteins that change subcellular location in the budding yeast *Saccharomyces cerevisiae* following chemically-induced DNA replication stress using methyl methanesulfonate (MMS) or hydroxyurea (HU). How these protein movements are regulated remains largely unexplored. We find that the essential checkpoint kinases Mec1 and Rad53 are responsible for regulating the subcellular localization of 159 proteins during MMS-induced replication stress. Unexpectedly, Rad53 regulation of the localization of 52 proteins is independent of its known kinase activator, Mec1, and in some scenarios independent of Tel1, or mediator components, Rad9 and Mrc1. We demonstrate that Rad53 is phosphorylated and active following MMS exposure in cells lacking Mec1 and Tel1. This non-canonical mode of Rad53 activation depends partly on the retrograde signaling transcription factor Rtg3, which also facilitates proper DNA replication dynamics. We conclude that there are biologically important modes of Rad53 protein kinase activation that respond to replication stress and operate in parallel to Mec1 and Tel1.

## Introduction

The DNA replication machinery serves to faithfully duplicate the genome in a highly regulated manner. Consistent with DNA replication being an essential cellular process, failure to replicate the genome leads to several physiological defects, including development disorders, neurodegeneration, growth defects, and predisposition to cancer (Aguilera and García-Muse, 2013; Ubhi and Brown, 2019; Zeman and Cimprich, 2014). Endogenous and exogenous forms of DNA damage can act as obstacles that impair the processivity of the replication fork machinery, resulting in replication stress. Chemically inducing methylation of nucleotide bases with methyl methanesulfonate (MMS) or depleting the nucleotide levels within a cell using hydroxyurea (HU) have both been shown to stall the replication fork (Pellicioli et al., 1999; Tercero et al., 2003). All forms of replication stress result in stretches of single-stranded DNA (ssDNA) that is rapidly coated with replication protein A complex (RPA), the signal for replication checkpoint activation.

The eukaryotic replication checkpoint serves to promote survival in conditions of replication stress by preventing cell-cycle progression and increasing nucleotide levels to allow for proper DNA repair. In the budding yeast, *Saccharomyces cerevisiae,* RPA-coated ssDNA is recognized and bound by Ddc2, which recruits its binding partner and an essential replication checkpoint kinase, Mec1 (Lisby et al., 2004; Pardo et al., 2017). Several protein factors at stressed forks, including the Rad17/Mec3/Ddc1 complex and Dpb11, activate and/or potentiate Mec1 kinase activity. Mec1 subsequently phosphorylates histone H2A at Ser129, promoting the recruitment of a mediator protein Rad9, which is then phosphorylated (Gilbert et al., 2001). Mec1 can also phosphorylate the checkpoint mediator and component of the replication fork complex (RFC) Mrc1, which causes dissociation from Pol ϵ and mediates Rad53 phosphorylation by Mec1 (Alcasabas et al., 2001; Chen et al., 2014; Zou and Elledge, 2003). Both mediators are capable of binding Rad53 to increase its local protein concentration, stimulating Rad53 trans-autophosphorylation and full activation (Branzei and Foiani, 2006; Wybenga-Groot et al., 2014). Hyperphosphorylated Rad53 then amplifies and transduces the signal cascade through several of its substrates, including another checkpoint kinase, Dun1 (Chen et al., 2007; Lee et al., 2003).

Numerous replication stress and DNA repair proteins organize into multiprotein complexes that can be observed as discrete nuclear foci using fluorescence microscopy (Gallina et al., 2015; Lisby et al., 2003; Tkach et al., 2012). This regulated re-localization of proteins is a key feature of the replication checkpoint response. Indeed, several proteins involved in Mec1 activation (Ddc1, Ddc2, and Dpb11) form subnuclear foci when cells are treated with MMS or HU (Bonilla et al., 2008; Lisby et al., 2004). In addition, the spatiotemporal organization of recombination repair proteins to foci has proven to be a valuable tool in dissecting the genetic requirements for the order of assembly of repair foci (Gallina et al., 2015; Tkach et al., 2012). Several studies have expanded on the repertoire of proteins that re-localize during replication stress by utilizing the budding yeast GFP collection to systematically probe protein subcellular location during replication stress conditions (Breker et al., 2013; Chong et al., 2015; Dénervaud et al., 2013; Mazumder et al., 2013; Tkach et al., 2012). We previously assessed protein location by manual inspection after treatment with HU or MMS and observed 254 proteins changing localization (Tkach et al., 2012). Interestingly, protein movement involved almost every cellular organelle, covering ten localization categories, with many localizations not involving the nucleus (Tkach et al., 2012). Finally, we, and others, have identified two distinct classes of nuclear foci. These include the DNA damage repair foci linked to DNA repair processes, and the intranuclear quality control compartments (INQ) involved in the turnover of replication stress factors, demonstrating that protein localization can provide insight into functional processes (Gallina et al., 2015; Tkach et al., 2012). While replication stress-induced protein localization has been well documented and has provided functional insight, the details of how these movements are regulated remains elusive.

Mec1 and Rad53 have a combined total of 1200 protein targets that they phosphorylate during replication stress and DNA damage conditions (Bastos de Oliveira et al., 2015; Chen et al., 2007; Lanz et al., 2018), and are therefore prime candidates for regulators of protein localization changes during replication stress. In this study, we investigated how Mec1 and Rad53 control protein localization dynamics. We employ a high-throughput confocal imaging platform to monitor subcellular location changes of 322 proteins known to re-localize during MMS or HU treatment. We find that checkpoint signalling is responsible for 159 (49%) of the protein localization changes observed. Our analysis also revealed a subset of proteins whose proper subcellular localization depended only on Rad53, and was independent of the known canonical upstream activators of Rad53. Finally, we demonstrate that Rtg3 promotes phosphorylation of Rad53 during replication stress, revealing a biological connection between retrograde signalling and the DNA replication stress checkpoint response.

## Results

### Checkpoint kinases Mec1 and Rad53 regulate replication-induced protein localization

Mec1 and Rad53 are two essential checkpoint kinases that initiate and propagate the replication checkpoint signal to effector proteins via phosphorylation, with Rad53 directly phosphorylated by Mec1 (Figure 1A). Previous studies have identified 322 proteins that change subcellular location following replication stress, re-localizing with varying dynamics following MMS and HU treatment (Chong et al., 2015; Dénervaud et al., 2013; Mazumder et al., 2013; Srikumar et al., 2013; Tkach et al., 2012; Table S1). Considering that these proteins are enriched for MMS-dependent phosphorylation targets, we assessed the role of the essential checkpoint kinases on their relocalization. We used Synthetic Genetic Array (SGA) to introduce *MEC1* or *RAD53* null mutations, in addition to an *SML1* deletion required for cell survival, into an array of 322 strains, each expressing one of the GFP-protein fusions previously found to re-localize during replication stress. The resulting strains were imaged by high-throughput confocal microscopy and protein localization was computationally quantified as previously described (Ho et al., 2022). Briefly, cells were grown to mid-logarithmic phase and imaged over four hours following 0.03% MMS treatment (Figure 1B). For each time point, the proportion of cells in the population with a protein-GFP localization change after MMS treatment was calculated (Table S2). To facilitate comparisons among the control and checkpoint mutants, the percent maximum value (% max) was calculated normalizing each timepoint to the maximum percent of cells for a given protein re-localization across the three screens (Figure 1C and D, Table S3). We also compared each mutant screen to the control screen by calculating log_2_ ratios of localization percentage values (% max) for each time point (Figure 1E). To reveal proteins with similar re-localization patterns, we clustered the resulting measurements for all three screens together using a *k*-means algorithm such that proteins with similar localization trajectories were grouped together. The analysis converged on seven *k*-means clusters (Figure 1C, D, and E).

**Figure 1:**
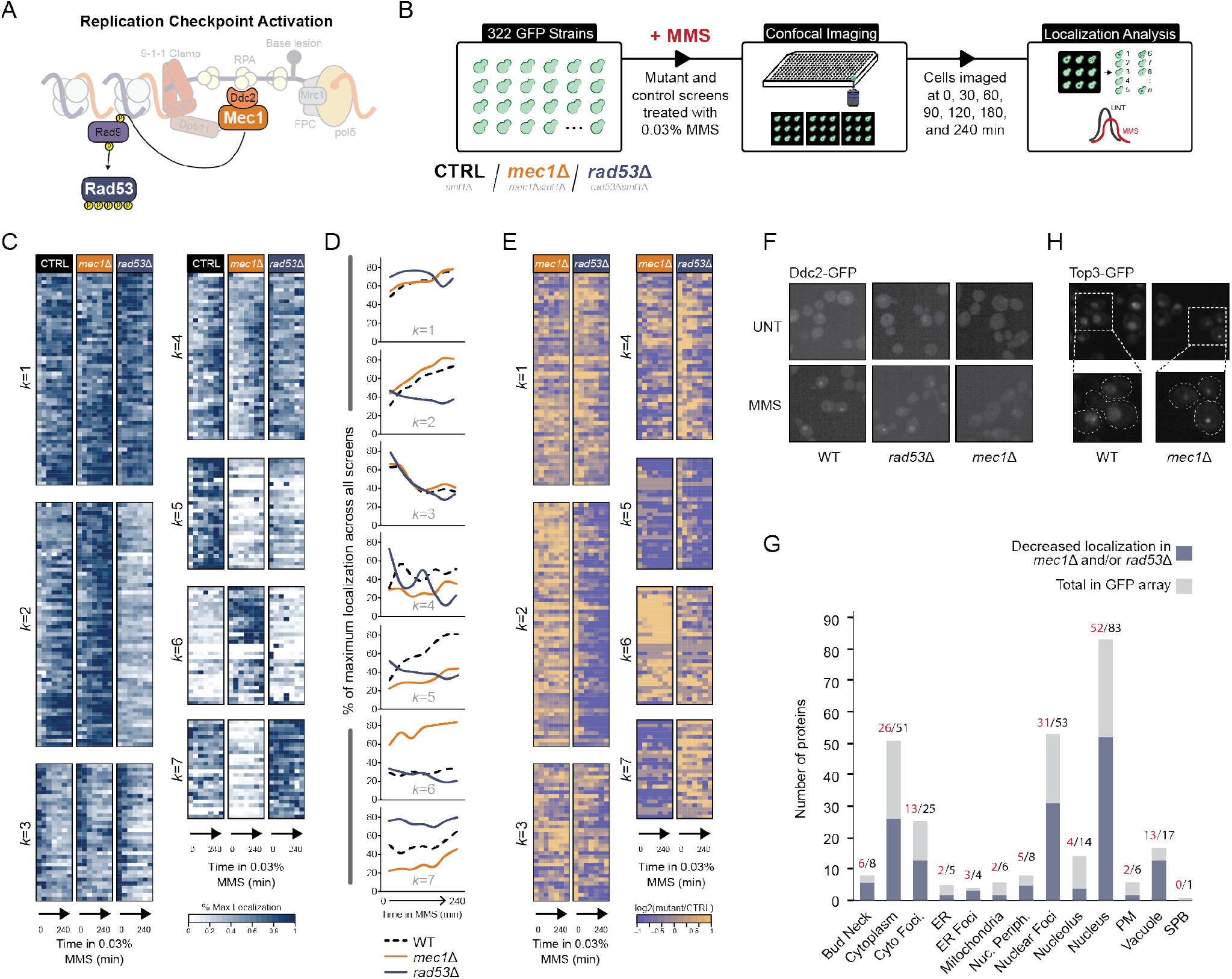
Checkpoint kinases Mec1 and Rad53 regulate global protein re-localization. (A) A simplified schematic of replication checkpoint activation in yeast. (B) Outline of the imaging and quantification pipeline used in this study. Briefly, *sml1*Δ, *mec1*Δ*sml1*Δ, or *rad53*Δ*sml1*Δ deletions were introduced into a set of 322 strains, each expressing a unique protein-GFP fusion. Mutant strains were treated with 0.03% MMS and imaged at 7 times over the course of 240 minutes. Resulting images were subsequently segmented, and cells with changes in protein localization were measured. (C) Heatmaps of the percent of maximum protein re-localization in MMS (% max) for the control, *MEC1,* and *RAD53* deletion screens. The resulting localization vectors were clustered into 7 groups by *k*-means, with the *k*-group number indicated to the left of the heatmap. (D) Line plots showing the mean extent of protein re-localization for each mutant screen at each time point for all proteins within respective *k*-group indicated. (E) The log_2_ ratio of the re-localization in mutant vs control is displayed for each timepoint. (F) Representative images of Ddc2-GFP nuclear focus formation in WT, *mec1*Δ, and *rad53*Δ cells after 2 hours treatment with 0.03% MMS. (G) Representative image of Top3-GFP nuclear foci in *mec1*Δ cells in unperturbed conditions. (H) The total number of proteins in each category of protein re-localization for all proteins assessed in our study is shown in grey. The number of proteins with evidence of decreased protein re-localization rates in either *mec1*Δ or *rad53*Δ cells is shown in blue.

Upon calculation of the average localization change over time within each *k*-cluster, distinct patterns were evident. The MMS-induced re-localizations of the 40 proteins in cluster *k*=5 depends on both *MEC1* and *RAD53* (Figure 1C, D, and E), indicating that those proteins are downstream of the replication stress signal transduced by Mec1 to Rad53 to target protein. The 39 proteins in cluster *k*=7 showed decreased re-localization only when *MEC1* was deleted, suggesting that these proteins are regulated by Mec1 but not by Rad53. As an example, the localization of Ddc2 to nuclear foci depends on *MEC1* but not on *RAD53* (Figure 1F), consistent with previous analysis indicating that Ddc2 localization is regulated by Mec1 and not by Rad53 (Melo et al., 2001). Two proteins (Rad50, Rfa1) in cluster *k*=7 were identified as Mec1 targets in phosphoproteomic analyses (Bastos de Oliveira et al., 2015), and four (Mms21, Msh3, Nej1, Rad50) contain SQ/TQ motifs, which are consensus sites for Mec1 phosphorylation. Thus, cluster *k*=7 contains proteins that are subject to Mec1 regulation, independent of Rad53. In total, 56% (159/284) proteins that change intracellular location (at least 2-fold change in the percentage of cells with re-localization compared to wildtype cells at the corresponding time point) in response to MMS-induced DNA replication stress are regulated, either directly or indirectly, by the checkpoint kinase cascade in the expected fashion: their re-localization either depends on *MEC1* alone, or depends on both *MEC1* and *RAD53.* We further noted that the proteins with reduced localization in checkpoint mutants are found in 12 of the 15 subcellular compartments represented in our study (Figure 1G). These data suggest that the replication checkpoint kinases regulate a global protein localization response following replication stress, with targets that are not solely restricted to the nuclear compartment.

We also noted that deletion of either kinase can result in proteins exhibiting increased re-localization, particularly in the absence of DNA replication stress (Figure 1C, D, and E). For example, Top3 nuclear focus formation is increased when *MEC1* was deleted (Figure 1H). The 36 proteins in cluster *k*=6, on average, show increased re-localization in *mec1*Δ even at t=0. Similarly, the 39 proteins in cluster *k*=7 show increased re-localization in *rad53*Δ that is evident in the absence of replication stress. A likely explanation for the increased spontaneous re-localization that we observe is that deletion of either kinase causes increased DNA replication stress, a phenomenon that has been documented extensively (Brush et al., 1996; Chen and Kolodner, 1999; Craven et al., 2002; Hoch et al., 2013; Motegi et al., 2006; Myung and Kolodner, 2002; Pennaneach and Kolodner, 2004; Shimada et al., 2002). Cells respond to genetically-induced replication stress by activating the checkpoint response in the absence of the stress induced chemically by MMS. Ninety-six proteins show increased re-localization at t=0 in a checkpoint kinase mutant, including a host of proteins that repair stressed and collapsed DNA replication forks, further emphasizing DNA replication stress is present in checkpoint mutants.

### Rad53 functions during replication stress when Mec1 is absent

An unexpected and striking category of re-localization involved the 52 proteins in cluster *k*=2 (Figure 1C), which displayed reduced re-localization when Rad53 is absent, but were largely unaffected by *mec1*Δ. Mec1 has long been recognized as the predominant upstream activator of Rad53. *In vitro* studies have unambiguously shown that Mec1 phosphorylates Rad53 and mediates the transautophosphorylation of Rad53 (Pellicioli et al., 1999; Sweeney et al., 2005). In the absence of Mec1, activation of Rad53 by replication stress or DNA damage is not detected by standard protein kinase assays or by the characteristic mobility shift on SDS-PAGE that accompanies Rad53 activation. Yet our data indicate that a large number of protein re-localization events depend on *RAD53*, and on the presence of DNA replication stress, but apparently not on *MEC1*.

One example of a *RAD53*-dependent, *MEC1* – independent re-localization is shown in Figure 2A. The recombination repair protein Rad54 forms nuclear foci in response to replication stress. The fraction of cells displaying Rad54 in nuclear foci decreases in *rad53*Δ cells, but not in *mec1*Δ cells (Figure 2A and B). In subsequent assays, we use Rad54 re-localization as an exemplar of Mec1-independent regulation by Rad53. We first established that decreased Rad54 re-localization in *rad53Δ* is not due to cell cycle effects or to Rad54 abundance changes.

**Figure 2:**
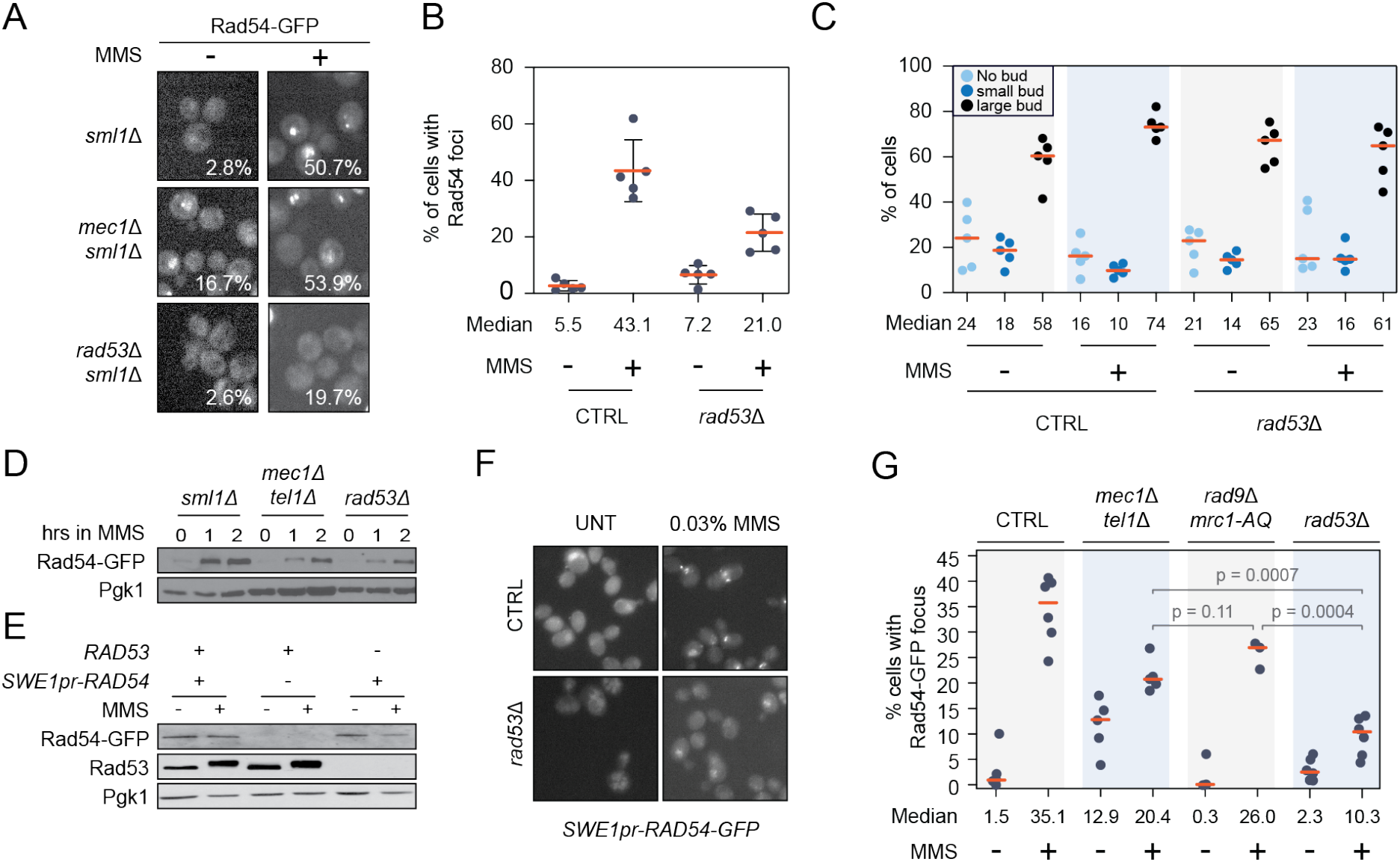
Rad54 nuclear foci formation requires Rad53 but not canonical replication checkpoint signaling. (A) Representative images of Rad54-GFP foci formation in *sml1*Δ, *mec1*Δ*sml1*Δ, or *rad53*Δ*sml1*Δ cells treated with 0.03% MMS for 2 hours. The percentage of cells with Rad54-GFP focus formation is indicated. (B) Quantification of the percentage of live cells with at least one Rad54-GFP nuclear focus in untreated and 2 hours of 0.03% MMS treatment for the indicated strains. The median percent of cells with Rad54-GFP foci is shown below (n=5). (C) The percentage of unbudded, small-budded, and large-budded cells in the indicated mutants, in untreated and MMS treatment. The median percent of cells for each category is shown at the bottom of the plot. (D, E) Mid-logarithmic phase cells for the indicated strains were fixed with TCA at the indicated times following 0.03% MMS exposure. Cells were lysed, and immunoblotted with anti-GFP, anti-Pgk1, or anti-Rad53 antibody. (F) Live cells were grown to mid-logarithmic phase, treated with MMS, and imaged by fluorescence microscopy (GFP) to observe Rad54-GFP focus formation. (G) Quantification of the percentage of live cells with at least one Rad54-GFP foci in each of the indicated strains, before and after 0.03% MMS treatment for 2 hours. The median percentage is shown at the bottom of the plot. For all plots, the orange horizontal bar indicates the median percentage of cells.

Cell cycle checkpoints arrest cells in G2-M in the presence of MMS to provide cells time to repair DNA damage (Kupiec and Simchen, 1985; Siede, 1995). Given that Rad53 is important for cell cycle arrest, it is formally possible that decreased Rad54 focus formation could be due to abnormal cell cycle progression in *rad53*Δ. It is well-documented that *rad53*Δ mutants progress more rapidly through S phase relative to wild type cells in conditions of replication stress (McClure and Diffley, 2021; Tercero et al., 2003). Importantly, the decrease in Rad54 foci in *rad53*Δ is clear at 2 hours, a time when both *rad53*Δ and wild type cells display similar proportions of large-budded cells (Figure 2C). Therefore, it is unlikely that the decrease in Rad54 foci is due to cell cycle effects.

Replication stress activates transcriptional and translational programs, ultimately modulating protein levels to facilitate DNA synthesis and repair (Jaehnig et al., 2013; Workman et al., 2006). We tested if Rad54 protein levels are reduced in *rad53*Δ cells compared to wild type after MMS treatment, which could also explain the decrease in nuclear foci. In the absence of *RAD53,* Rad54 protein levels are similar to wild type cells (Figure 2D). Rad54 protein levels increased after MMS treatment, and the increase was greater in wild type cells than in *mec1*Δ*tel1*Δ or *rad53*Δ cells. However, Rad54 levels were similar in *mec1*Δ*tel1*Δ and *rad53*Δ cells, suggesting that the different levels of Rad54 foci in *mec1*Δ*tel1*Δ and *rad53*Δ cells are not an effect of altered Rad54 abundance (Figure 2D). Additionally, when we replaced the *RAD54* promoter with the *SWE1* promoter the cells no longer displayed an increase in Rad54 protein abundance after MMS exposure (Figure 2E), but still formed Rad54 foci in the presence, but not in the absence, of *RAD53* (Figure 2F). We conclude that Rad54 focus formation is not strictly linked to Rad54 abundance.

Another potential explanation for the absence of *MEC1* dependence for Rad54 re-localization is that yeast carry a *MEC1* orthologue, *TEL1,* that in some instances can provide functional redundancy in activating Rad53 (De la Torre-Ruiz et al., 1998; Morrow et al., 1995; Vialard et al., 1998). We asked whether Rad53 still promotes Rad54 focus formation in cells with null mutations in both *MEC1* and *TEL1.* Cells expressing Rad54-GFP from the native *RAD54* locus were treated with 0.03% MMS for two hours and Rad54 nuclear focus formation was measured. We detected a decrease in Rad54 focus formation in *mec1Δtel1Δ* compared to wild type cells, but deletion of *RAD53* caused a greater decrease, indicating that in the absence of *MEC1* and *TEL1, RAD53* makes independent contributions to Rad54 re-localization (Figure 2G). Assessing Rad54 foci in *mec1*Δ *tel1*Δ *sml1Δ* was confounded by severe growth defects and morphological abnormalities, both of which could lead to an underestimate of re-localization. As an alternative approach, we mutated *MRC1* and *RAD9,* which encode the checkpoint mediators that propagate the replication stress signal from Mec1/Tel1 to Rad53 (Figure 2G). In the absence of mediator function the re-localization of Rad54 is unaffected, consistent with Rad54 re-localization being independent of signalling from Mec1/Tel1 to Rad53. Together, our data suggest that Rad54 focus formation depends on a pathway that transduces the signal elicited by MMS treatment to the checkpoint kinase Rad53 without a requirement for Mec1 and Tel1.

### Rad53 kinase activity and function is required for Rad54 focus formation

We next considered the domains of Rad53 that could promote protein relocalization without activation by Mec1/Tel1 and the mediators Mrc1 and Rad9. Since canonical activation of Rad53 was not essential for Rad54 re-localization, it was possible that a kinase-independent function of Rad53 was involved. In addition to its kinase domain, Rad53 contains two forkhead-associated domains at its N- and C-terminal ends, termed FHA1 and FHA2 (Bashkirov et al., 2003; Durocher et al., 1999; Liao et al., 1999). FHA1 functions in binding some Rad53 substrates, and FHA2 is important for interaction with Rad9 and subsequent Rad53 activation by trans-autophosphorylation (Durocher et al., 1999, 2000; Liao et al., 1999; Matthews et al., 2014; Pike et al., 2003). We quantified the percentage of cells with Rad54-GFP nuclear foci in cells expressing inactivating mutations in each of the domains (R70A, R605A, and K227A/D319A/D339A (KD), located in the FHA1, FHA2, and kinase domains, respectively). Only the inactivation of the kinase domain *(RAD53-KD)* resulted in decreased Rad54 re-localization to the level seen in *rad53Δ* cells (Figure 3A). Interestingly, mutation of FHA2 had only a modest effect, again consistent with the canonical Mec1-mediator axis of Rad53 activation playing at best a minor role in Rad54 re-localization. These results indicate that Rad53 kinase activity is necessary for Rad54 re-localization in response to DNA replication stress, whereas neither FHA domain plays a substantial role.

**Figure 3:**
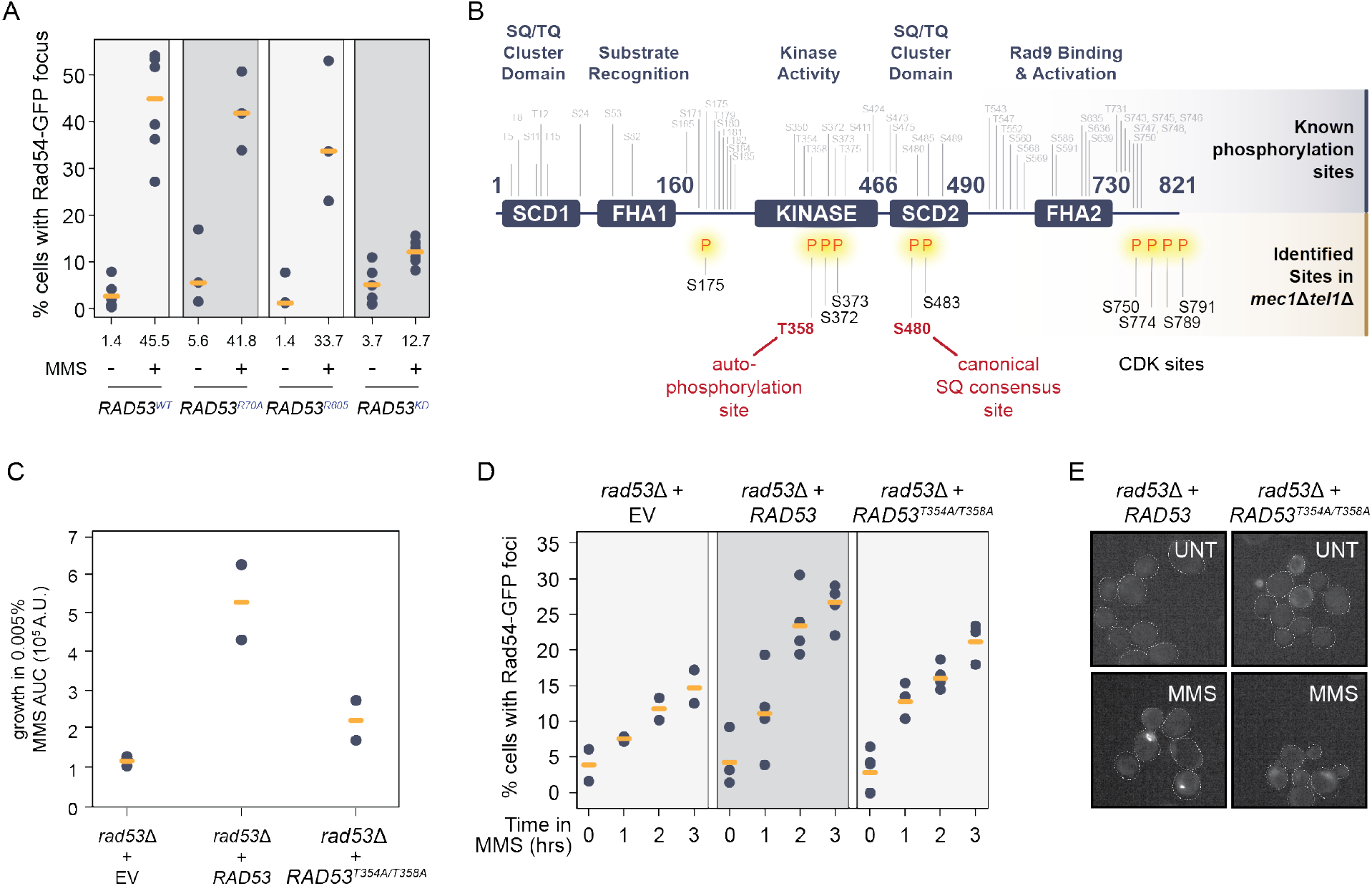
Rad53 is phosphorylated independent of Mec1/Tel1 and its kinase activity promotes Rad54 re-localization. (A) Quantification of the percentage of live cells with at least one Rad54-GFP foci in each of the indicated strains, before and after 0.03% MMS treatment for 2 hours. The median percentages are indicated by the orange bars, and shown for each. (B) Schematic of the domain architecture of the Rad53 protein, with known phosphorylation sites annotated above in grey. Phosphorylation sites in *mec1*Δ*tel1*Δ cells identified by mass spectrometry are annotated below the schematic in black, with two highlighted in red representing functional phosphorylation sites described in the literature. (C) The growth of cells with the indicated mutations was measured and the growth of cells was estimated by the area under the curve. (D) Quantification of the percentage of live cells with at least one Rad54-GFP foci in the indicated Rad53 mutants over three hours of 0.03% MMS treatment. (E) Representative images of Rad54-GFP foci in *rad53*Δ cells expressing the indicated *RAD53* alleles.

Rad53 is normally activated following priming phosphorylation by Mec1, which promotes Rad53 dimerization and trans-autophosphorylation (Gilbert et al., 2001; Ma and Stern, 2008; Pellicioli and Foiani, 2005; Wybenga-Groot et al., 2014). Given that Rad53 kinase activity is tightly coupled to its phosphorylation, and that we find evidence for Rad53 activity outside of the characterized Mec1 pathway, we tested whether Rad53 was phosphorylated in *mec1*Δ *tel1*Δ cells. Rad53 fused with a 10X-FLAG epitope was expressed from its native locus, and cells were treated with MMS. Following affinity purification and phosphopeptide enrichment, Rad53 phosphopeptides were identified by mass spectrometry (Figure 3B, Table S4). We found only 10 phosphorylated residues in *mec1*Δ*tel1*Δ cells, which is in stark contrast to the 48 sites on Rad53 normally occupied by phosphorylation in DNA damage conditions. Unexpectedly, phosphorylation was detected within the kinase and SCD2 domains of Rad53. Serine 480 is a Mec1 S/T-Q consensus site in the SCD2 domain, and promotes Rad53 activity when phosphorylated (Lee et al., 2003). Our data show that S480 can be phosphorylated by a kinase distinct from Mec1 and Tel1. Of particular interest, we detected phosphorylation of T358, which is a transautophosphorylation site important for full activation of Rad53 (Fiorani et al., 2008; Ma and Stern, 2008). Phosphorylation of T358 suggests that Rad53 dimerization and trans-autophosphorylation have occurred, resulting in Rad53 kinase activation. We mutated T358 and the adjacent autophosphorylation site at T354 to alanine *(RAD53-TA* mutant) and found that it sensitized cells to MMS, and reduced Rad54 re-localization (Figure 3C, D, and E). Taken together, these results indicate that Rad53 can be phosphorylated in response to DNA replication stress in a Mec1- and Tel1-independent manner, and explain the important role of the Rad53 kinase activity in Rad54 re-localization despite the lack of a strong requirement for Mec1/Tel1 or mediator proteins.

### Retrograde signaling factor Rtg3 promotes Mec1-independent Rad54 focus formation

Our data suggest that Rad53 can respond to MMS, undergo dimerization, autophosphorylate, and promote proper localization of Rad54, all independent of Mec1 and Tel1 function. Therefore, we sought to identify the molecular factors involved in this non-canonical mode of Rad53 activation. Given that the replication checkpoint is a phosphorylation signaling cascade, and Rad53 itself is phosphorylated in MMS, we focused on protein kinases and kinase-related genes. We catalogued genes annotated in the *Saccharomyces Genome* Database (https://www.yeastgenome.org/) as either possessing or promoting kinase or phosphatase activity. We generated an array of 296 strains, each expressing Rad54-GFP from the *RAD54* locus and a unique gene deletion or temperature sensitive allele. To focus on Mec1-independent Rad53 activity, we also deleted *MEC1* in these strains (Figure 4A). We measured the percentage of cells with a change in Rad54 localization after 2 hours of MMS treatment (Figure 4B, Table S5). We identified 31 gene deletion mutants that had decreased Rad54 foci (p < 0.05, student’s t-test). The top three genes identified in our screen that resulted in reduced Rad54 focus formation are all part of the retrograde (RTG) signalling pathway *(RTG2, RTG3, and MKS1).* The RTG pathway regulates the expression of genes involved in metabolism and mitochondrial maintenance (Jazwinski and Kriete, 2012; Jia et al., 1997; Komeili et al., 2000; Liao and Butow, 1993; Ruiz-Roig et al., 2012), but has no known role in checkpoint signalling. To validate the screen results, we reconstructed *rtg2*Δ *mec1*Δ and *rtg3*Δ *mec1*Δ mutants, and assessed Rad54 focus formation induced by MMS (Figure 4C and D). Both strains displayed reduced levels of Rad54 re-localization compared to the respective single mutants. In particular, *rtg2*Δ*mec1*Δ had strikingly low levels of Rad54 foci, similar to *rad53*Δ cells, indicating that the RTG pathway influences Mec1-independent Rad53 function.

**Figure 4:**
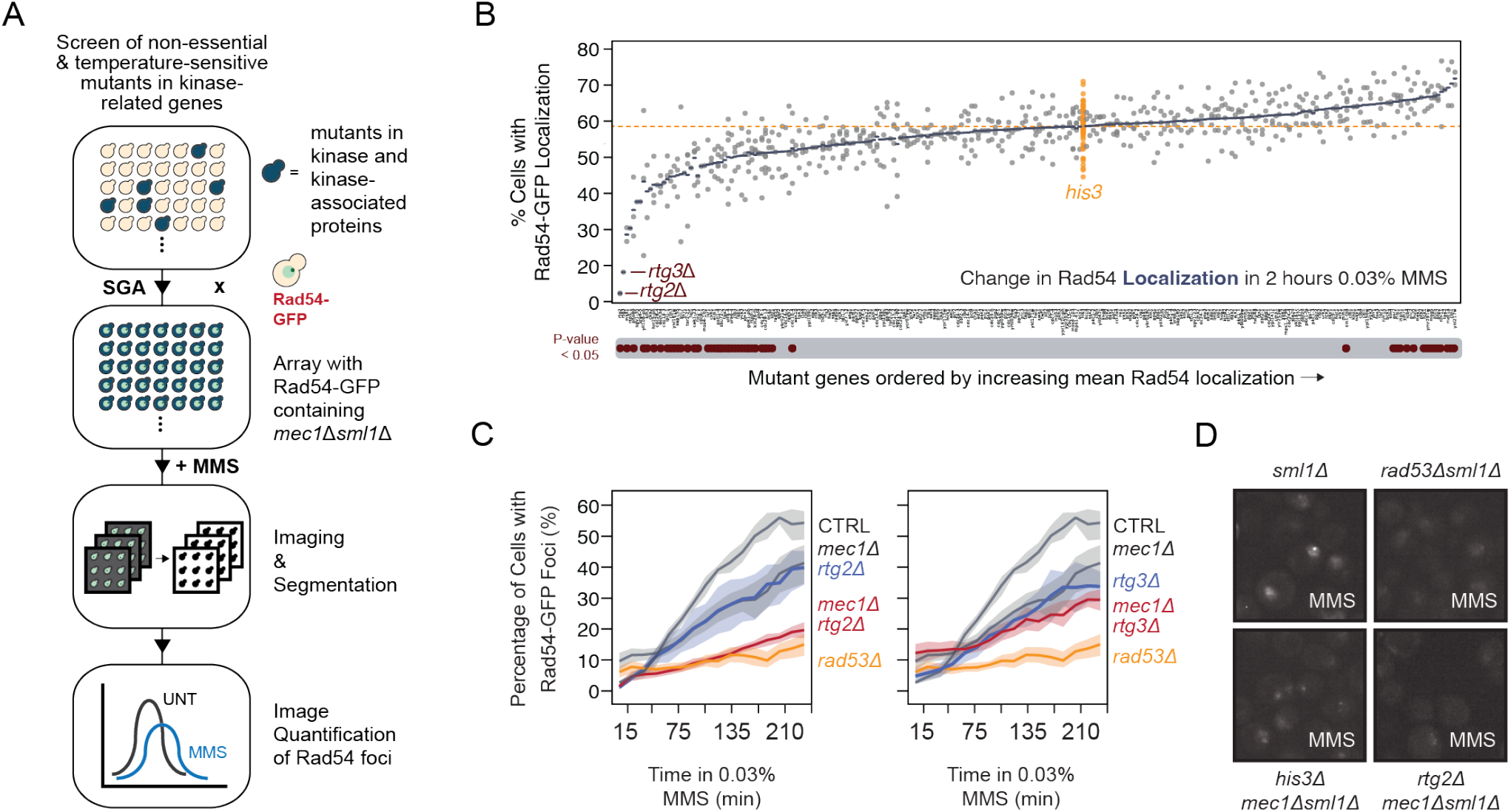
Retrograde signaling in promotes Rad54 foci formation in *mec1*Δ. (A) Outline of the imaging and quantification pipeline for the small scale screen. Briefly, *mec1*Δ*sml1*Δ cells expressing Rad54-GFP from its native locus were introduced by SGA into an array of unique strains each with a mutation in a kinase-related gene. The resulting strains were treated with MMS, and imaged on a high throughput confocal microscope. (B) The percentage of cells with a change in Rad54-GFP localization was determined for all kinase-related genes, indicated at the bottom of the plot. The percent of cells with Rad54-GFP re-localization for 60 independent control *his3*Δ mutants are shown in the orange points. Genes with reduced levels of foci compared to the mean of all *his3*Δ deletion control mutants are indicated below the plot (p < 0.05). For reference, the orange horizontal dotted line represents the mean of all *his3*Δ controls. (C) Quantification of percent of cells with Rad54-GFP foci after treatment with 0.03% MMS for the indicated times for the *rtg2*Δ and *rtg3*Δ mutants identified from the screen (n = 3). (D) Representative images of Rad54-GFP foci in the indicated mutant strains.

### The retrograde signalling pathway responds to replication stress

If the RTG pathway is transducing a replication stress signal to Rad53, we reasoned that the RTG pathway should respond to replication stress. The heterodimeric Rtg1/Rtg3 transcription factor is the downstream effector of the RTG pathway (Figure 5A) (Workman et al., 2006) and translocates from the cytoplasm to the nucleus when the pathway is activated (Sekito et al., 2000), so we focused on *RTG3.* Microscopic inspection of Rtg3-GFP revealed that Rtg3 translocates to the nucleus following MMS-induced replication stress, and persists in the nucleus for longer when the MMS concentration is increased (Figure 5B). Therefore, Rtg3 responds in a quantifiable manner to DNA replication stress, a prerequisite if Rtg3 transmits a replication stress signal to Rad53.

**Figure 5:**
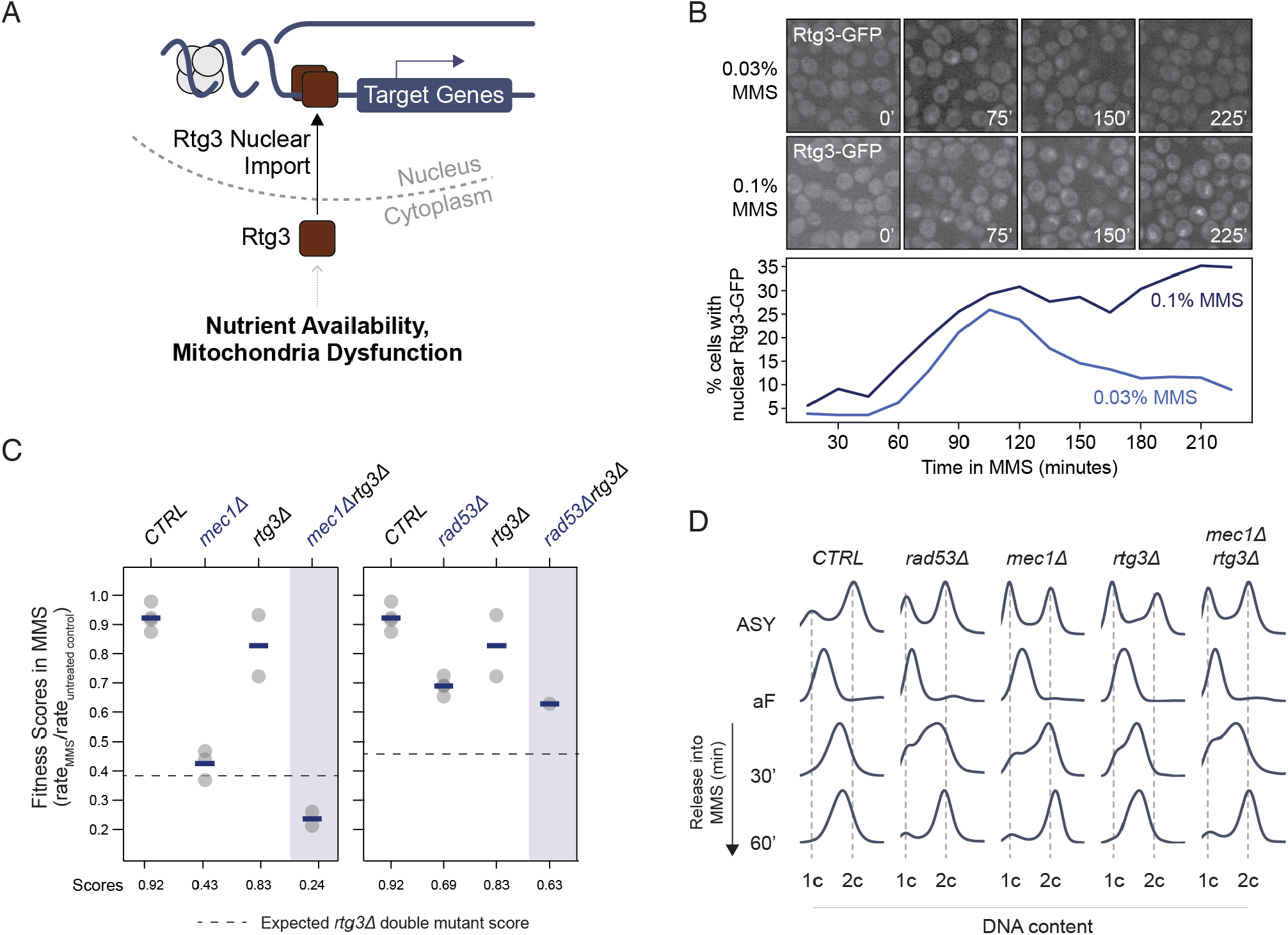
Nuclear accumulation of Rtg3 is important for regulating genes involved in stress responses. (A) Under conditions of nutrient limitation or mitochondrial dysfunction, Rtg3 enters the nucleus and upregulates stress-responsive genes. (B) Representative confocal micrographs and quantification of the percent of live cells with Rtg3-GFP nuclear localization after 0.03% MMS treatment for 4 hours. (C) Fitness scores for the indicated deletion mutants grown in 0.005% YPD liquid culture were calculated by growth rate in MMS / growth rate in untreated conditions. *RTG3* genetic interactions were assessed in (left) *mec1*Δ or (right) *rad53*Δ cells, and indicated in the dashed lines are the expected double mutant scores calculated according to the multiplicative model. A negative genetic interaction would be a score that falls below this line. (D) Cell cycle analysis of the indicate strains. Cells were arrested in G1 with 250ug/mL alpha-factor for 2.5 hours, then released into 0.03% MMS. Samples were fixed and prepared for DNA content analysis by flow cytometry at the indicated time points. Positions of 1C and 2C content are indicated.

Deletion of *RTG1* or *RTG3* confers sensitivity to DNA replication stress induced by hydroxyurea (Hartman IV, 2007), and so we tested if deletion of *RTG3* confers MMS sensitivity, and whether *RTG3* is in the same replication stress response pathway as *RAD53*. The *rtg3*Δ strain showed a small fitness defect in MMS (Figure 5C). When we combined *rtg3*Δ with *mec1*Δ, the fitness defect in MMS was greater than expected from analysis of the single mutants. Importantly, when we combined *rtg3*Δ with *rad53*Δ, no greater-than-additive or greater-than-multiplicative fitness defect was evident, indicating that *RTG3* functions in MMS resistance in the same genetic pathway as *RAD53,* and parallel to *MEC1* (Figure 5C).

Since the checkpoint kinases are important in proper cell cycle progression and replication fork stabilization during replication stress (Bermejo et al., 2011; Iyer and Rhind, 2013; De Piccoli et al., 2012; Toledo et al., 2017), we sought to directly assess S-phase progression in *rtg3*Δ cells. Cells were arrested in G1 phase and released synchronously into S phase in the presence of MMS (Figure 5D), where the peak is shifted further left compared to the control at 30 minutes). As expected, *mec1*Δ and *rad53*Δ cells exhibited S phase defects, both progressing faster than wild type cells and reaching 2C DNA content by 60 minutes (Shimada et al., 2002), whereas wild type cells remain in S phase with less than 2C DNA contents. At 30 minutes, we noted that *rad53*Δ cells (where the peak of DNA contents remained less than 2C) showed a slower S-phase progression compared to *mec1*Δ cells (where the peak of DNA contents reached 2C). The S phase progression of *rtg3*Δ *mec1*Δ cells closely resembled that of *rad53*Δ cells, indicating that *RTG3* influences the replication stress response when *MEC1* is absent (Figure 5D). We conclude that *RTG3* responds to DNAreplication stress, promotes resistance to replication stress, and functions within the *RAD53* pathway, parallel to *MEC1*.

### Rad53 phosphorylation is reduced when Rtg3 signaling is removed

If the RTG pathway is promoting signalling via Rad53 in the absence of Mec1, then we would expect that the activating phosphorylations that we identified on Rad53 (Figure 3B) might depend on Rtg3 function. We immunoprecipitated Rad53-FLAG in *rtg3*Δ *mec1*Δ and *mec1*Δ cells, and used quantitative mass spectrometric analysis to assess the Rad53 phosphorylation status in these mutants (Figure S1). Strikingly, we found six sites that were reduced by at least two-fold in *rtg3*Δ*mec1*Δ compared to *mec1*Δ (S469, S774, S473, S424, S766, and S743; Figure 6A, Table S6). An additional four sites were reduced by at least 1.5-fold (T563, S745, S489, and S475). Notably, three of these sites (S473, S480, and S489) are Mec1/Tel1 consensus phosphorylation sites, and S480 and S489 are located within the Rad53 SCD2 domain, which is important for Rad53 oligomerization and activation (Ma and Stern, 2008). Perhaps surprisingly, we did not detect changes in phosphorylation at T358. One possible reason may be the presence of redundant activities that promote T358 phosphorylation. Nevertheless, our data provide concrete evidence that the RTG pathway promotes phosphorylation of Rad53 on activation-relevant sites, in the absence of Mec1, consistent with the RTG pathway providing an unrecognized mode of Rad53 activation that can function in parallel to canonical Mec1-dependent Rad53 activation.

**Figure 6:**
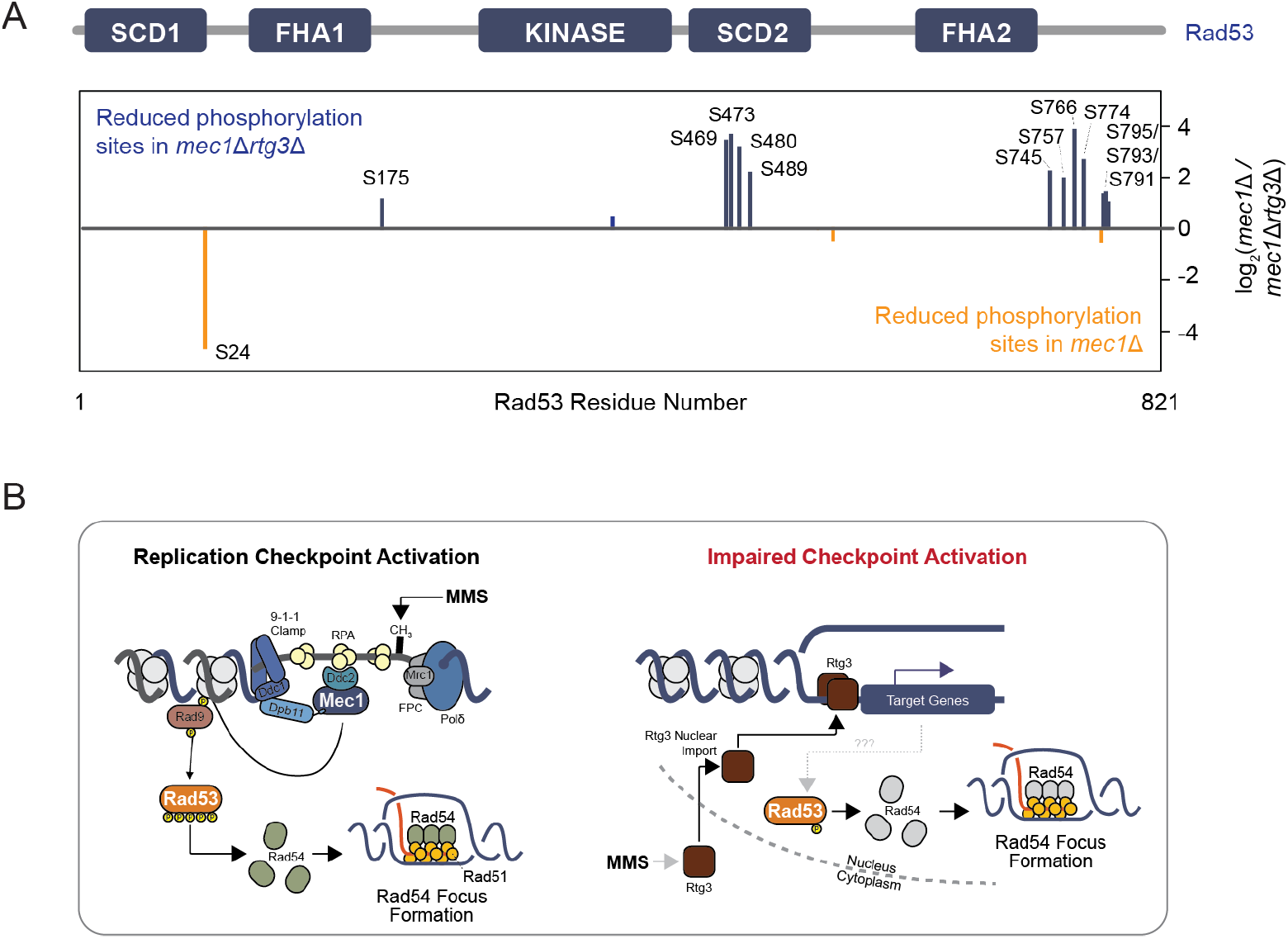
Rad53 phosphorylation is reduced when Rtg3 signaling is removed. (A) Rad53-FLAG was immunoprecipitated with anti-flag in *mec1*Δ*sml1*Δ cells with or without *rtg3*Δ, phosphopeptides were enriched, and quantified by SILAC-MS. Phosphopeptides that are enriched or depleted in *mec1*Δ*sml1*Δ*rtg3*Δ cells are indicated in orange and blue, respectively. A schematic of the Rad53 protein and its domains shows the location of the differentially phosphorylated sites. (B) A model for Rad53 activation in DNA replication stress when the canonical checkpoint signaling axis is impaired *mec1*Δ cells. Replication stress signals for the nuclear import of the Rtg3 transcription factor, resulting in gene expression changes. Potential target genes of Rtg3 promote Rad53 phosphorylation at functionally important residues, stimulating limited Rad53 activity, which is required for proper Rad54 focus formation.

## Discussion

How the replication checkpoint regulates changes in protein subcellular location is largely unknown. By measuring replication stress induced protein re-localizations in *mec1*Δ and *rad53*Δ cells, we showed that the check-point kinases coordinate a global protein re-localization response and regulate the movement of 159 proteins during replication stress. Most importantly, the proper localization of 52 proteins depend on Rad53, but not on the upstream kinases or mediator proteins of the replication checkpoint signaling cascade. We show that the RTG signalling pathway contributes to this novel mode of Rad53 activation. We propose a model (Figure 6B) where cells with an impaired replication checkpoint (i.e. *mec1*Δ cells) sense and respond to MMS-induced damage via the RTG pathway, resulting in nuclear accumulation of Rtg3. Since Rtg3 is a transcription factor, we postulate that Rtg3 regulates the expression of a kinase or kinases capable of phosphorylating Rad53. Alternatively, Rtg3 may function as a scaffold that promotes Rad53 dimerization and subsequent phosphorylation. The resulting phosphorylation of Rad53 appears to sustain at least a subset of Rad53 functions. How the DNA replication stress is sensed, and the additional effectors that function in this pathway, remain to be discovered.

### Rad53 functions independent of canonical replication checkpoint signaling

Even though the replication checkpoint has been studied extensively, we suggest that Rad53 can be activated in the absence of its known activators, Mec1 and Tel1. Consistent with our inference, independent lines of evidence support the hypothesis that Rad53 exhibits genome stability functions independent of Mec1 and Tel1. In unperturbed cell cycles, *rad53*Δ cells display slower S-phase progression than *mec1*Δ (Manfrini et al., 2012), an effect that we also observed during replication stress (Figure 5D). Rad53 is recruited to replication fork proximal sites independent of the mediator Mrc1, suggestive of checkpoint-independent Rad53 recruitment (Sheu et al., 2021). Rad53 also binds to gene promoters independently of Mrc1 and Mec1, again indicating Rad53 function independent of Mec1 (Sheu et al., 2021). *MEC1* and *RAD53* have genetically distinct roles in stabilizing stressed replication forks (Lanz et al., 2018; Segurado and Diffley, 2008), although Mec1-independent function of Rad53 was not noted. Finally, Rad53 function in the degradation of excess histones is Mec1-independent (Gunjan and Verreault, 2003). We demonstrated that Rad53 is phosphorylated in MMS-induced DNA replication stress in cells lacking Mec1 and Tel1. Two sites were of particular interest: (1) a canonical S/T-Q phosphorylation site at Ser480 in the SCD2 domain, known to be important for Rad53 oligomerization, and (2) an autophosphorylation site at Thr358 that, when mutated to alanine, severely compromises full Rad53 phosphorylation (Chen et al., 2014; Ma and Stern, 2008; Wybenga-Groot et al., 2014). Our data are consistent with the idea that Rad53 kinase contributes to replication stress resistance even in the absence of its main activators, Mec1 and Tel1.

### Mec1-independent activation of Rad53 occurs through dimerization and phosphorylation

The typical mode for Rad53 activation requires its phosphorylation-dependent interaction with checkpoint mediator proteins Mrc1 or Rad9 (Durocher et al., 2000; Sweeney et al., 2005). Upon replication stress, Mec1 is recruited to and activated at RPA-coated ssDNA, where it then phosphorylates Rad9, among many other targets (Rouse and Jackson, 2002; Zou and Elledge, 2003). Mec1 phosphorylation enables Rad9 binding to the Rad53 FHA2 domain. Mec1 then phosphorylates several residues in the Rad53 SCD1 domain, resulting in the oligomerization, autophosphorylation, and full activation of Rad53 (Sanchez et al., 1999). Our indicates that Mec1-independent activation of Rad53 occurs in a similar manner. The phosphorylation at Ser480 of Rad53 that we detect in the absence of Mec1 promotes oligomerization, and phosphorylation at Thr358 of Rad53 in the absence of Mec1 indicates that dimerization and trans-autophosphorylation has occurred. Since Rad9 and Mrc1 are not required for the Mec1-independent Rad53 activity that we observe, we propose that additional factors serve to increase the local concentration of Rad53 enough to promote oligomerization and transautophosphorylation. Concentration-dependent activation of Rad53 has been observed upon overexpression of Rad53 in bacteria (Gilbert et al., 2001), consistent with local concentration of Rad53 being sufficient to cause at least partial activation. Therefore, it is reasonable to speculate that a portion of total Rad53 protein can locally concentrate, activate, and exert its functions, independent of the canonical checkpoint activating factors Mec1, Tel1, Rad9, and Mrc1.

### Rtg3 functions to maintain genome stability

Changes in the mitochondrial state, or its dysfunction, activates a transcriptional program that adjusts metabolic activities and stress responses and is termed retrograde (RTG) regulation (Giannattasio et al., 2005; Jazwinski, 2013). In yeast, Rtg1 and Rtg3 are basic helix-loop-helix leucine zipper transcriptional activators that translocate into the nucleus when cells experience mitochondrial dysfunction. Rtg2 is another key protein component of the RTG signaling pathway and is required for Rtg1/3 nuclear accumulation. RTG signaling integrates with another signaling pathway that senses intracellular stress, the TOR pathway (Crespo et al., 2002). Unexpectedly, we found that RTG signaling is also involved in genome maintenance. Most strikingly, our finding that Rtg3 contributes to Rad53 phosphorylation status implicates retrograde response factors in replication checkpoint signalling, Rad53 function, and cellular fitness upon exposure to DNA replication stress.

Several lines of evidence support the notion that retrograde signaling impinges upon genome integrity pathways. Rtg2 inhibits the formation of extrachromosomal ribosomal DNA circles (ERC), a form of genome instability (Borghouts et al., 2004). ERCs arise from recombination events between distal sites within a chromosome and are self-replicating episomes of the ribosomal DNA (Stults et al., 2008; Szostak and Wu, 1980). Accumulation of ERCs has been correlated with yeast ageing, and circular DNAs capable of driving genetic heterogeneity and therapeutic resistance are abundant in certain human cancer cells (Koche et al., 2019; Yan et al., 2020). Our understanding of circular DNA biogenesis, regulation, and function is far from complete. Nevertheless, the connection between Rtg2 and ERCs suggest roles for retrograde signaling in maintenance of genome integrity.

Rtg3 also cooperates with Ino4 to regulate the expression of metabolism genes after MMS exposure (Workman et al., 2006). TCA cycle regulation, threonine biosynthesis, and permease trafficking pathways coordinate with one another to buffer dNTP pools, suggesting an indirect role of RTG in maintaining proper cellular nucleotide levels, which are crucial for normal DNA replication (Koche et al., 2019). Importantly, *rtg1, rtg3, and rtg3* deletion mutants exhibit severe growth defects in conditions of nucleotide limitation and have reduced levels of cellular dNTPs. In contrast, *RTG3* overexpression induces ribonucleotide reductase expression, further linking *RTG3* to dNTP pool regulation (our unpublished data). Consistent with a role in replication, Rtg3 was identified in a genome wide search for factors that facilitate replication of long inter-origin gaps (Theis et al., 2010). Similarly, we found that Rtg3 promotes proper S-phase progression, in a pathway that is dependent on Rad53 kinase activity yet does not require Mec1 signaling. Like RTG signaling, Rad53 has essential roles in elevating dNTP levels for repair, preventing late origin firing, and phosphorylating Mrc1 to antagonize CMG helicase unwinding and slow the replication fork (Lopez-Mosqueda et al., 2010; McClure and Diffley, 2021; Morafraile et al., 2015, 2019; Szyjka et al., 2008; Tsaponina et al., 2011). Since our data shows that Rtg3 influences Rad53 phosphorylation status and function, we suggest that *RTG3* functions in the completion of DNA replication during replication stress, at least in part through Rad53 activation. Thus, in addition to identifying the complement of proteins whose location is regulated by cell cycle checkpoint kinases, we have found a previously uncharacterized mode of checkpoint signalling.

## Methods Details

### Yeast Strains and Media

All yeast strains were in the S288C (BY) background. Unless otherwise stated all strains with either a *MEC1* or *RAD53* deletion also contain an *SML1* deletion. Strains were constructed using genetic crosses and standard PCR-based gene disruption and epitope tagging techniques.

### Generating checkpoint mutant GFP collections by synthetic genetic array

An array comprised of 322 yeast strains was created, each consisting of unique ORF tagged with GFP and the following genotype: *MAT***a** *XXX-GFP::his3MX his3*Δ1 *leu2*Δ0 *met15*Δ0 *ura3*Δ0. Three query strains were constructed, which include (1) *mec1*Δ::kanMX *sml1*Δ::hphMX, (2) *rad53*Δ::natMX *sml1*Δ::hphMX, and (3) *sml1*Δ::hphMX, each with the genotype *MATα can1pr-RPL39pr-tdTomato::CaURA3::can1*Δ::*STE2pr-LEU2 his3*Δ1 *leu2*Δ0 *lyp1*Δ0. Synthetic genetic array was performed as described (Baryshnikova et al., 2010; Tong et al., 2001) to introduce the gene deletions from the query strains into the GFP array of 322 strains. The final strains were selected on solid agar media SD/MSG – his – leu -lys – arg + 50 μg/mL canavanine + 50 μg/mL thialysine + 300 μg/mL hygromycin. The *MEC1* and *RAD53* deletion strains were selected on the same media with the addition of 160 μg/mL geneticin and 100 μg/mL nourseothricin, respectively.

### High-throughput confocal microscopy

For protein localization screens involving the 322 GFP mini array, cells were grown to logarithmic phase at 30° C and imaged by GFP fluorescence microscopy using the OPERA High Content Screening System (PerkinElmer), as described (Torres and Brown, 2015). Briefly, cells were grown in low fluorescence media (LFM – 1.7g/L yeast nitrogenous base without amino acids and without ammonium sulfate, 2% glucose, 1x methionine, 1x uracil, 1x histidine, and 1x leucine) in 96-well polypropylene U-bottom culture boxes to logarithmic phase at 30°C. Cells were then diluted to an OD600 = 0.02-0.05 in 384-well, clear bottom plates for microscopic imaging (PerkinElmer 6007550) and incubated at 30° C for 1h to allow cells to settle. The cells were imaged on the PerkinElmer OPERA imaging system before and after MMS treatment with the following configuration: primary dichroic filters reflecting excitation wavelengths of 488 nm and 561 nm, detection dichroic set to 568 nm, 520/35 nm emission filter for Camera 1, 600/35 nm emission filter for Camera 2, laser power set to 100%, and exposure for GFP and RFP channels at 800 ms.

For protein localization screens involving the kinase-related array, and GFP fluorescence assays involving Rad54-GFP imaged on the PerkinElmer OPERA Phenix System. Cells were grown in synthetic complete (SC) media (6.7g/L yeast nitrogenous base without amino acids and with ammonium sulfate, 2% w/v glucose, 1x amino acids) at 30°C to mid-logarithmic phase (OD_600_ = 0.3-0.7). Cells were diluted and prepared as above, treated with 0.03% MMS for the indicated timpoints, then imaged on the OPERA Phenix with the following configuration: primary dichroic filters reflecting excitation wavelengths of 488 nm and 561 nm, detection dichroic set to 568 nm, 520/35 nm emission filter for Camera 1, 600/35 nm emission filter for Camera 2, laser power set to 100%, and exposure for GFP and RFP channels at 800 ms.

### Calculating percentages of cells exhibiting a localization

For all images acquired from the OPERA and OPERA Phenix, image segmentation of single yeast cells was conducted using CellProfiler 3.1 (McQuin et al., 2018), GFP pixel intensity measurements of the resulting segmented single cells was performed using Python 3.7.0 (https://www.python.org), and data analyses and visualization were performed using Python and/or R (https://www.r-project.org). Briefly, for every cell in our screens, the 95th percentile of the GFP pixel intensity was calculated and normalized to the cell median GFP intensity, representing the “localization value” of a cell. For each unique protein-GFP fusion, the localization value was determined for untreated cells, and the median and median absolute deviation (MAD) for this distribution were calculated. For every timepoint before and after drug treatment, a cell with a localization value greater than or less than 1.5 MADs from the median was considered to a cell exhibiting a protein re-localization event. Calculation of the percent max value was performed as describe in (Ho et al., 2022). For the kinase-associated gene screen with cells expressing Rad54-GFP, the 98th percentile of GFP pixel intensity was used.

### Budding index and live cell GFP fluorescence microscopy

Cells expressing Rad54-GFP were grown in synthetic complete (SC) media (6.7g/L yeast nitrogenous base without amino acids and with ammonium sulfate, 2% glucose, and all amino acids supplemented) at 30°C to mid-logarithmic phase (OD_600_ = 0.3-0.7). Cells were either left unperturbed or treated with 0.03% MMS for 120 minutes. Cells were imaged on a Nikon Eclipse Series Ti-2 inverted widefield microscope using open software *μ* manager (https://micro-manager.org/). Budding index and the percentage of cells with a localization signal were manually assessed by visual inspection.

### Flow cytometry

For cell cycle analyses, 1mL of cultures at OD_600_ = 0.5 were collected at the indicated times, fixed in 70% ethanol, and stored at 4°C until sample processing. Cells were washed in ddH2O, resuspended in 0.5mL of 50mM Tris-Cl (pH 8.0) with 2 mg/mL RNase A, and incubated for 2 hr at 37°C. Cells were then pelleted, and resuspended in 0.5 mL of 50mM Tris-Cl (pH 7.5) with 1 mg/mL proteinase K (BioShop PRK403) and incubated for 1 hr at 50°C. Cells were pelleted, and resuspended in 200 mM Tris-Cl (pH 7.5), 200 mM NaCl, and 78 mM MgCl2, and stored at 4°C. Immediately before analyzing samples on the flow cytometer, 0.1 mL of sample was added to 0.5 mL SYBR green solution (1:5000 dilution, Sigma-Aldrich S9430), sonicated briefly, and analyzed using a FACS Canto II (Becton Dickinson). A total of 10 000 events were collected, and plots were generated using FlowJo software version 10.0.8.

### Rad53 phosphorylation mapping

Yeast expressing RAD53-FLAG tagged at its native locus were grown to mid-log phase in YEPD medium and treated with 0.03% methyl methansulfonate for 2 hours. Cells were lysed by bead-beating with 0.5mm glass beads for 3 cycles of 10 minutes with a 1 minute rest between cycles at 4°C in lysis buffer (150mM NaCl, 50mM Tris pH 7.5, 5mM EDTA, 0.2% NP40) supplemented with complete EDTA-free protease inhibitor cocktail (Roche), 5 mM NaF and 10 mM β-glycerophosphate. RAD53-FLAG was immunoprecipitated from approximately 5mg of protein extract with FLAG antibody-conjugated agarose resin and eluted with 1% SDS in 100mM Tris pH 8.0. Eluate was reduced using 10mM DTT and subsequently alkylated with 25mM iodoacetamide followed by precipitation for 1hr in PPT solution (50% acetone, 49.9% ethanol, 0.1% acetic acid) on ice. Pellets were washed once with PPT solution and then resuspended in urea/tris solution (8M urea, 50mM Tris pH 8.0). 8M urea containing solubilized pellet was diluted to 2M urea using milliQ water and then digested overnight at 37°C with 1ug of trypsin GOLD (Promega). Phosphopeptides were selectively enriched using a home-made Fe-NTA resin micro-column and then subjected to LC-MS/MS analysis on a Thermo-Fisher Q-Exactive HF mass spectrometer. Raw MS/MS spectra were searched using SORCERER (Sage N Research, Inc.) running SEQUEST software over a composite *S. cerevisiae* peptide database consisting of normal protein sequences downloaded from the Saccharomyces Genome Database (SGD) and their reversed protein sequences as a decoy to estimate the false discovery rate (FDR) in the search results. Searching parameters included a semi-tryptic requirement, a mass accuracy of 15 ppm for the precursor ions, a static mass modification of 57.021465 daltons for alkylated cysteine residues, and a differential mass modification of 79.966331 daltons for phospho-serine, threonine and tyrosine residues.

### Whole cell extracts, co-immunoprecipitation, and immunoblotting

Cells were cultured and grown to OD_600_ = 0.3-0.5 and treated with 0.03% MMS for the indicated times. Cells were then diluted to OD_600_ = 1.0 in 10% trichloroacetic acid for 15 minutes, shaking gently, then pelleted by centrifugation at 3000 rpm for 5 minutes. Next, cells were resuspended in 2X Laemmli sample buffer and lysed by vortex on maximum speed in the presence of 0.5 mm glass beads, at 4°C for 10 minutes. Samples were boiled at 95°C for 10 minutes, and stored long term at −80°C. Proteins were resolved on SDS-PAGE gels and detected by immunoblotting with anti-Rad53 (1:5000, Abcam ab104232), anti-PGK (1:1 000 000, Novex 459250), anti-GFP (1:5000, ClonTech 632375), and anti-FLAG M2 (1:5000, Sigma F3165).

### Growth rate assays

Saturated cultures of yeast strains were diluted to an OD_600_ = 0.05 in 100 μL of YPD (with or without the indicated concentrations of drug) in flat bottom 96-well plates. Plates were inserted into a TECAN microplate analyzer set at 30° C with gentle agitation and the OD600 was measured every 10 minutes for 48 hours. The growth rate (maximum slope) and the area under the curve (AUC) was calculated using an R package designed by Danielle Carpenter (https://scholar.princeton.edu/sites/default/files/botsteinlab/files/growth-rate-using-r.pdf).

### Computational Analyses, Data, and Software Availability

Statistical analysis, data manipulation, and data visualization were performed in R (https://www.r-project.org) or Python (https://www.python.org). All the details of the data analysis can be found in the Results and Methods sections. Python and R scripts used for data processing, analysis, and visualization are available online on GitHub (https://github.com/bqho). Complete raw image data available from the authors upon request.

## Acknowledgements

This work was supported by the Canadian Institutes for Health Research (FDN-159913 to GWB), the National Institutes of Health grant (R35GM141159 to M.B.S.), an Ontario Government Scholarship, and a natural Sciences and Engineering Research Council of Canada CGS-D award (to B.H.). GWB holds a Canada Research Chair (Tier 1). We thank Raphael Loll-Krippleber for helpful discussions and careful reading of the manuscript. We are grateful to work on the lands of the Mississaugas of the Credit, the Anishnaabeg, the Haudenosaunee and the Wendat peoples, land that is now home to many diverse First Nations, Inuit, and Métis peoples.

## Author Contributions

Conceptualization, B.H., N.P.T., and G.W.B.; Methodology, B.H., N.P.T., R.L.K., and G.W.B.; High-throughput screening, B.H. and N.P.T.; Experimentation; B.H.; Massspectrometry, E.S. and M.B.S.; Formal Analysis, B.H., E.S., M.B.S., and G.W.B.; Writing – Review & Editing, B.H., E.S., M.B.S., and G.W.B.

---

**Figure S1:**
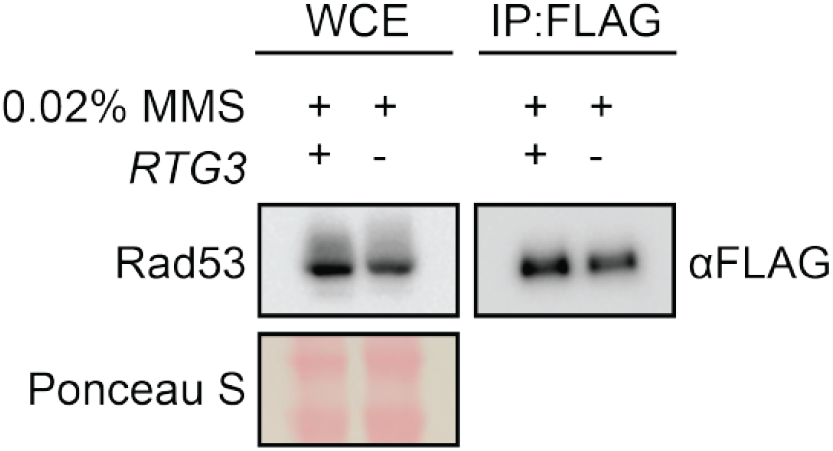
Immunoprecipitation of Rad53-FLAG for SILAC-MS. The levels of Rad53-FLAG immunoprecipitated for SILAC-MS analyses in *mec1*Δ *sml1*Δ cells with or without *rtg3*Δ was observed by immunoblotting with anti-flag.

